# Effects of Beaver (*Castor canadensis*) Herbivory and Wildfire on Foliage Density and Woody Debris, San Pedro Riparian National Conservation Area, Arizona

**DOI:** 10.1101/2022.06.25.497364

**Authors:** Marcia F. Radke

## Abstract

Beaver (*Castor canadensis*) were reintroduced beginning in 1999 on the San Pedro Riparian National Conservation Area. Herbivory is the most obvious effect from beaver, but little research exists of herbivory effects after reintroduction. Fire processes may also have substantial effects to vegetation, and both beaver and fire are important ecological components for comparing effects and subsequent management decisions. The objective of this research, conducted during 2008 and 2009, was to determine any effects to foliage density caused by beaver herbivory and wildfire as compared to control sites. There were significant differences in foliage density between control, beaver, and wildfire sites, with lower foliage density and greater above-ground heights associated with wildfire sites. Although near the significance level, there were no interactions between control, beaver, or wildfire sites for changes in foliage density at different heights. Mean Fremont cottonwood, Goodding’s willow, and seep willow foliage density was significantly different between control, beaver, and fire sites. Fremont cottonwood had significantly higher foliage density at control sites than at fire sites, but not between control and beaver sites or between beaver and fire sites. Goodding’s willow density was significantly higher at control and beaver sites than fire sites, with no significant difference between control and beaver sites. Seep willow foliage density was significantly higher at control and beaver sites compared to fire sites, but not significantly different between control and beaver sites. Mean downed and dead wood cover was not significantly different between control, beaver, and fire-influenced sites, between beaver and control sites, between control and fire sites, or between beaver and fire sites. Management implications include more strategic wildfire planning and preparedness, achieved through integrated tamarisk control in the riparian area and use of prescribed fire in upland habitats to reduce fire size and severity that threaten the riparian gallery forest and its ecosystem services.

## INTRODUCTION

After extirpation by fur trappers by 1894 (Bailey 1971), beaver (*Castor canadensis*) were reintroduced on the San Pedro Riparian National Conservation Area (SPRNCA) during 1999, 2000, and 2002 in a coordinated effort between Bureau of Land Management (BLM) and Arizona Game and Fish Department. Based on an average of 5.2 beaver per colony (Collen and Gibson 2001), and about 20 colonies with 33 dams, the estimated beaver population on the SPRNCA was 100 by 2008.

In other areas, beaver herbivory has shown to be an important component in shaping vegetation communities, including tree density and basal area (Johnston and Naiman 1990). The effects of beaver and avian community structure have been studied on the San Pedro River (Johnson and van Riper 2014), but the effect of beaver herbivory to vegetation has not been previously studied on the SPRNCA. Effects to vegetation are important because of the occurrence of federally listed species that may nest in riparian vegetation, including southwestern willow flycatcher (*Empidonax traillii extimus*) and yellow-billed cuckoo (*Coccyzus americanus*). This riparian ecosystem is also an important habitat corridor for numerous wildlife species including other neotropical birds (Stromberg and Tellman 2009).

Similar to beaver herbivory, fire may also have discernable effects to relative cover (Busch 1995) and structure (Bendix and Cowell 2010) of riparian vegetation. Thus, it is important to document the effects of both beaver herbivory and wildfire compared to control areas, especially where beaver reintroduction has occurred. Foliage density is a clearly observable and quantifiable object affected by beaver herbivory and fire, and the purpose of this research was to quantify the effects to foliage density caused by beaver herbivory or wildfire compared to control sites.

## STUDY AREA

The SPRNCA, located in southeastern Arizona approximately 85-km (53-mi) southeast of Tucson, was established in 1988 with Public Law 100-696. This Congressional designation established the conservation area shall be managed “in a manner that conserves, protects, and enhances the riparian area…” by the BLM. The riparian area is the “river of green,” largely surrounded by Chihuahuan desert scrub, and contains 82-km (51-mi) of the San Pedro River immediately north of Sonora, Mexico. The river flows from its headwaters in Mexico north approximately 209-km (130-mi) to join the Gila River. Fremont cottonwood (*Populus fremontii*)/Goodding’s willow (*Salix gooddingii*) gallery forest occurs over the river’s length from the International Boundary to approximately the historic ghost town of Contention about 64 river km (40 mi) north of Mexico. Thereafter, the cottonwood/willow gallery forest continues, but is invaded by increasing amounts of non-native tamarisk (*Tamarix ramosissima*) to the SPRNCA’s northern boundary near St. David, Arizona.

## METHODS

Foliage density data was collected during 2008 to 2009 during summer months when plant species were completely leafed. Areas influenced by beaver herbivory were located using UTM coordinates of 2008 active dams, and only included those sites with active and pre-existing beaver herbivory where beavers were known to have occupied for the longest time period since reintroduction. Data collected at control, beaver, and wildfire sites included identification of species (Fremont cottonwood, Goodding’s willow, or seep willow – *Baccharis salicifolia*). Control sites were randomly chosen in the same river reach as beaver sites in order to limit variability with geomorphology, water regime, and vegetation differences. Wildfire sites were chosen using known wildfires that occurred in the riparian area on the SPRNCA.

Foliage density was estimated at 0 to 100% (in 5% segments) using a 1-m (3.3-ft) square density board (Sanders and Flett 1989). A total of 40 plots (20 each on both the west and east side of the river) at each site were read at each of 0 to 1, 1 to 2, and 2 to 3-m (0 to 3.3, 3.3 to 6.6 and 6.6 to 9.8-ft) heights at 5-m (16.4-ft) intervals. At beaver-influenced sites, 20 plots were read on the west side of the river, and 20 plots were read on the east side of the river, with the reader beginning at the dam. All plots were read by one observer. The species comprising the majority of the foliage in front of the board was the species recorded, and species recorded included Fremont cottonwood, Goodding’s willow, and seep willow. Down and dead wood density was recorded using the same method.

The significance threshold for all analyses was 0.05. The *a priori* design to test for differences in percent foliage density was a parametric three-factor analysis of variance (ANOVA) using influence (control, beaver, and wildfire as three levels), species (Fremont cottonwood, Goodding’s willow, seep willow, and wood as four levels, and 0 to 1, 1 to 2, and 2 to 3-m (0 to 3.3, 3.3 to 6.6, and 6.6 to 9.8-ft) heights as three levels. After data collection, I tested for normality for n≥20 using the D’Agostino-Pearson K^2^ test for normality (Zar 1999). Foliage density data sets were significantly different from normal; therefore, data lumping was utilized to increase the power of analyses when no significant differences were found within a factor. Species were lumped, and percent foliage density was analyzed using a two-factor ANOVA with treatment and height as the factors.

Assumptions for a parametric ANOVA were not met if a significant difference from normality existed for each data set. Significant differences in normality were exhibited with Fremont cottonwood, seep willow, and wood densities, but not for Goodding’s willow. Other data sets that were not normal included the difference in foliage density between Fremont cottonwood, seep willow, and woody debris. In this case, I used a nonparametric Kruskal-Wallis ANOVA with tied ranks to assess significance, and any significant difference between groups was determined using nonparametric Mann-Whitney pairwise comparisons. Foliage density for Goodding’s willow was analyzed using a one-way ANOVA with treatment (control, beaver, or wildfire-influenced) as the factor, and pairwise comparisons after ANOVA were analyzed using the Tukey test (Zar 1999).

## RESULTS

There was a significant difference in foliage density (with all species lumped) between control, beaver, and wildfire sites (Figure 1; two-way ANOVA, F=6.938, df=∞, P=0.001). Pairwise comparison indicated significant differences in foliage density between wildfire and control sites (Tukey test, q=5.135, df=∞, P=0.001) and between wildfire and beaver sites (Tukey test, q=4.676, df=∞, P=0.003), but not between control and beaver sites (Tukey test, q=0.566, df =∞, P=0.915). Lower foliage density was directed at wildfire sites.

**Figure 1.**
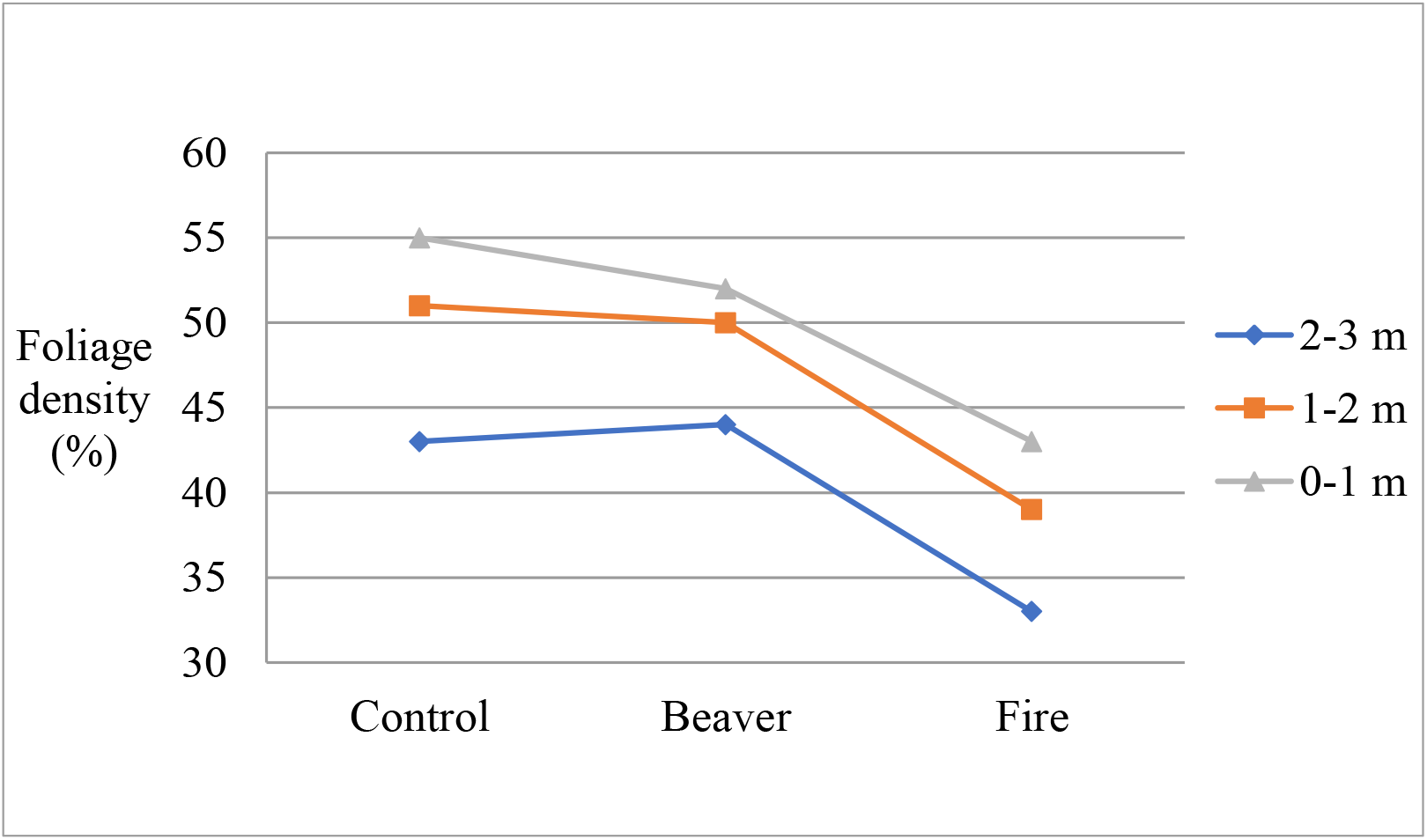
Mean foliage density (percent) at 0-1, 1-2, and 2-3 meter heights between control, beaver, and wildfire-influenced sites, San Pedro Riparian National Conservation Area, 2010.

Significant differences existed in foliage density at different heights (Figure 1; two-way ANOVA, F=7.544, df=∞, P=0.001). There was no significant interaction between control, beaver, or wildfire sites and changes in foliage density at different foliage heights (Figure 1; two-way ANOVA, F= 0.192, df = ∞, P=0.05). Significant differences in foliage density were found between control, beaver, and wildfire between 1 to 2 and 2 to 3 m (3.3 to 6.6 and 6.6 to 9.8 ft) heights (Tukey test, q=3.693, df=∞, P=0.025; Figure 1), and between 0 to 1 and 2 to 3 m (0 to 3.3 and 6.6 to 9.8 ft) heights (Tukey test, q= 5.374, df=∞, P=0; Figure 1). No significant difference existed between 0 to 1 and 1 to 2 m (0 to 3.3 and 3.3 to 6.6 ft) heights (Tukey test, q=1.732, df=∞, P=0.439; Figure 1). Lower foliage density was directed at increased height. Significant differences were found with five comparisons between foliage density by influence and height, with lower foliage density directed at wildfire sites and increased height (least significant Tukey test, q=4.392, df=∞, P=0.05). No significant difference existed between all other comparisons (most significant Tukey test, q=3.646, df=∞, P=0.197).

Mean Fremont cottonwood cover was significantly different between control, beaver, and wildfire sites (Figure 2; Kruskall-Wallis test, F=14.027, df=∞, P<0.001), with significantly higher foliage density at control sites than at wildfire sites (Mann-Whitney pairwise comparison, P=0.018). Mean foliage density of Fremont cottonwood was not significantly different between control and beaver sites (Mann-Whitney pairwise comparison, P=0.138) or between beaver and wildfire sites (Mann-Whitney pairwise comparison, P=0.243).

**Figure 2.**
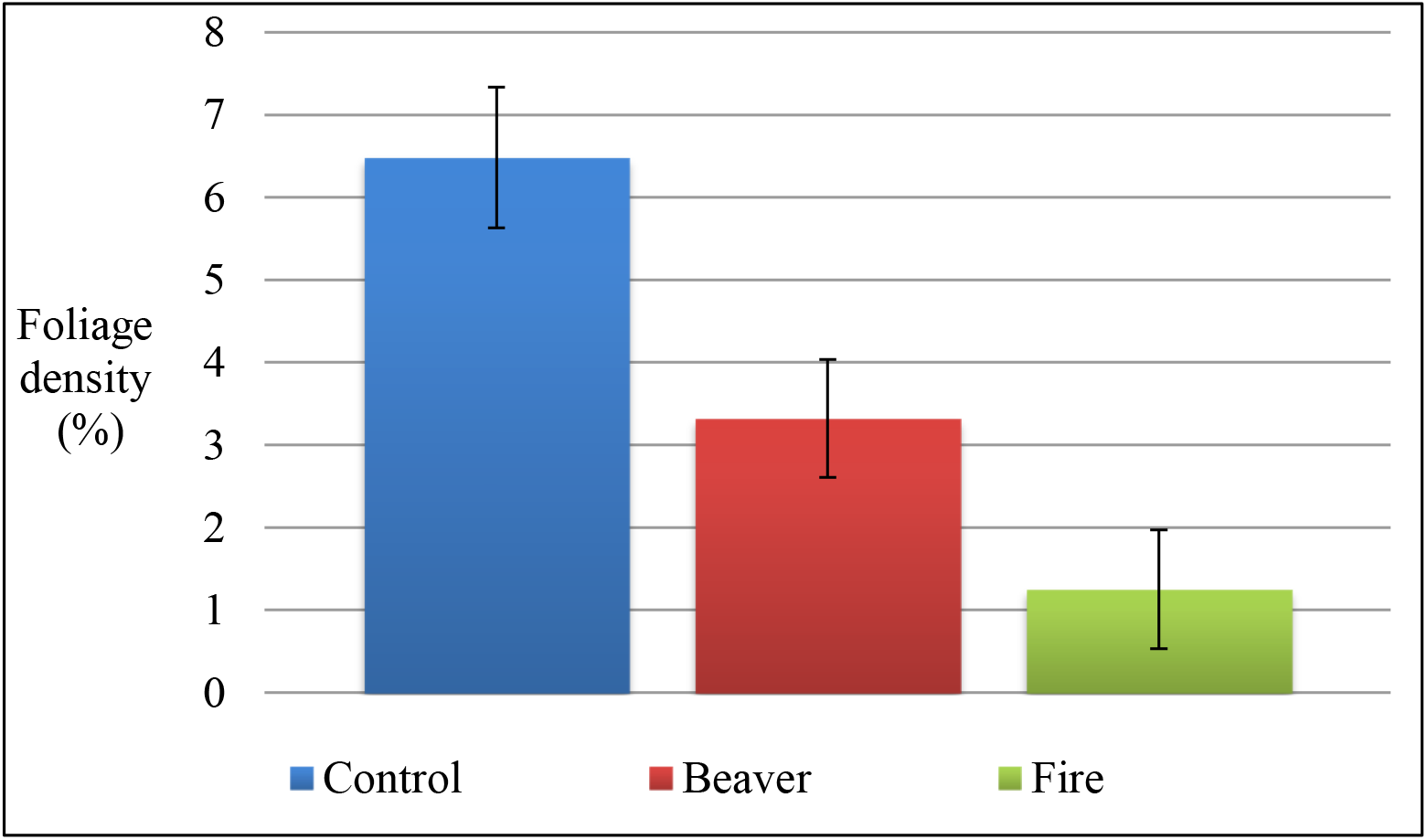
Mean (± standard error) foliage density of Fremont cottonwood at 0 to 3 m (0 to 9.8 ft) height between control, beaver, and wildfire sites, San Pedro Riparian National Conservation Area, 2008 to 2009.

Mean foliage density of Goodding’s willow cover was significantly different between control, beaver, and wildfire sites (Figure 3; one-way ANOVA, F=9.927, df=121, P=0), with higher foliage density at control sites (Tukey test, q=5.783, df=121, P=0) and at beaver sites (Tukey test, q=5.682, df=121, P=0) than at wildfire sites. Mean foliage density of Goodding’s willow was not significantly different between control and beaver sites (Tukey test, q=0.646, df=121, P=0.891).

**Figure 3.**
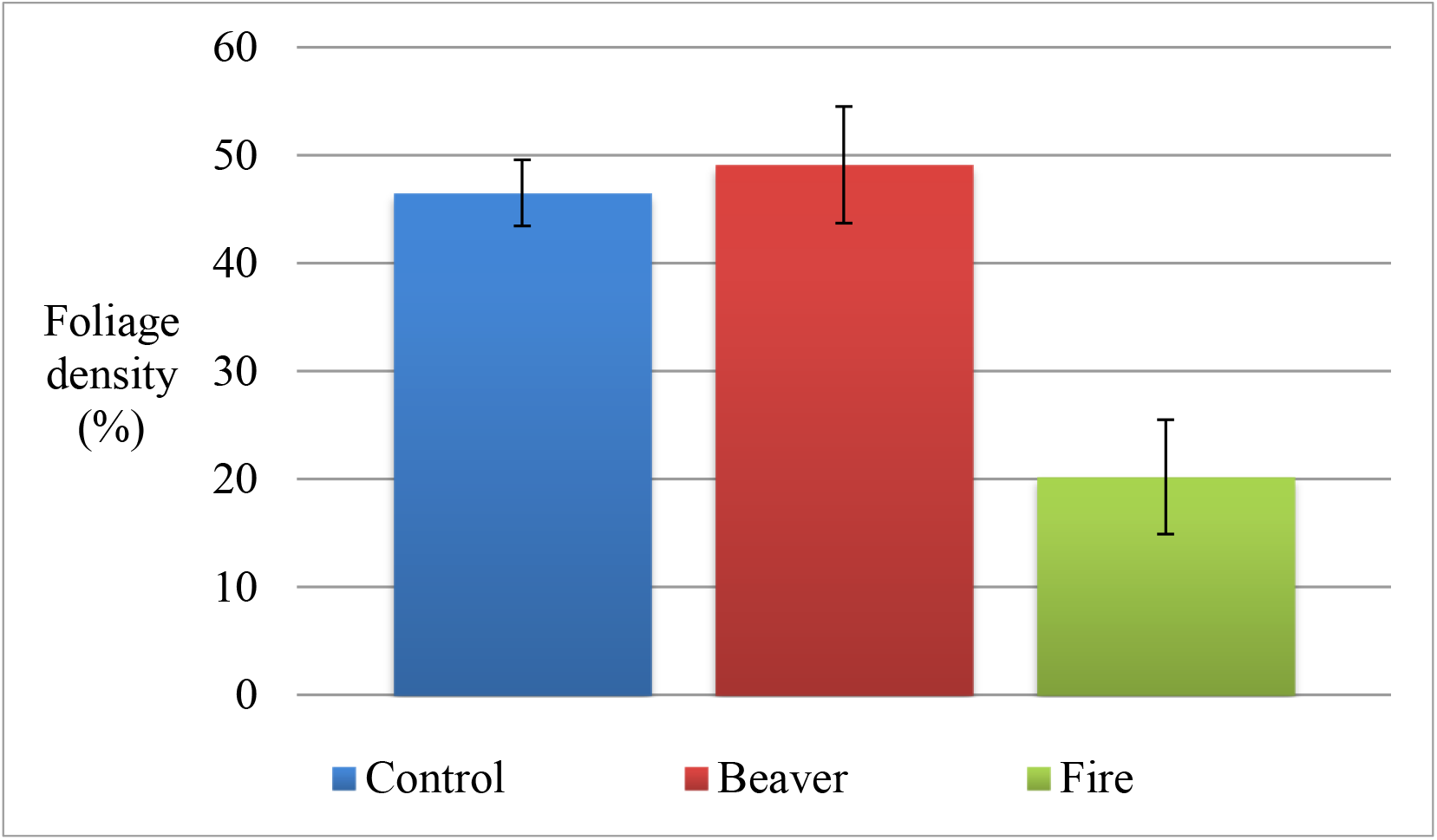
Mean (±standard error) foliage density of Goodding’s willow at 0-3 meter height between control, beaver, and wildfire-influenced sites, San Pedro Riparian National Conservation Area, 2010.

Mean seep willow cover was significantly different between control, beaver, and wildfire sites (Figure 4; Kruskall-Wallis test, F=4.185, df=∞, 0.02<P<0.05), with significantly higher foliage density at control sites (Mann-Whitney pairwise comparison, P=0.004) and beaver sites (Mann-Whitney pairwise comparison, P=0.02) than at wildfire sites. Mean foliage density of seep willow was not significantly different between control and beaver sites (Mann-Whitney pairwise comparison, P=0.71).

**Figure 4.**
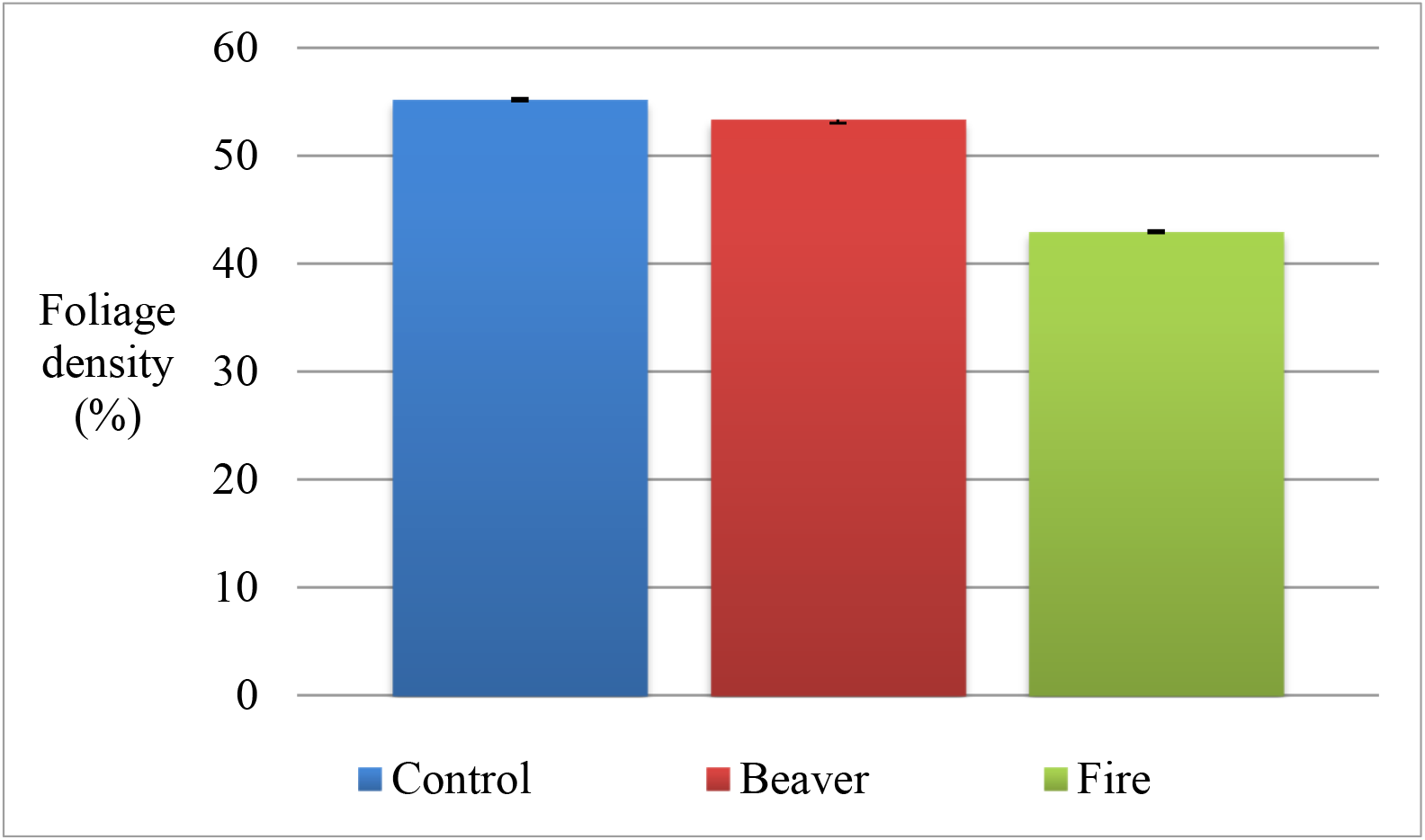
Mean (±standard error) foliage density of seep willow at 0-3 meter height between control, beaver, and wildfire-influenced sites, San Pedro Riparian National Conservation Area, 2010.

Mean woody debris cover was not significantly different between control, beaver, and wildfire sites (Figure 5; Kruskall-Wallis test, F=1.079, df=75, P>0.50), with no significant difference between beaver and control sites (Mann-Whitney pairwise comparison, P=0.15.), between control and wildfire sites (Mann-Whitney pairwise comparison, P=0.90), or between beaver and wildfire sites (Mann-Whitney pairwise comparison, P=0.29).

**Figure 5.**
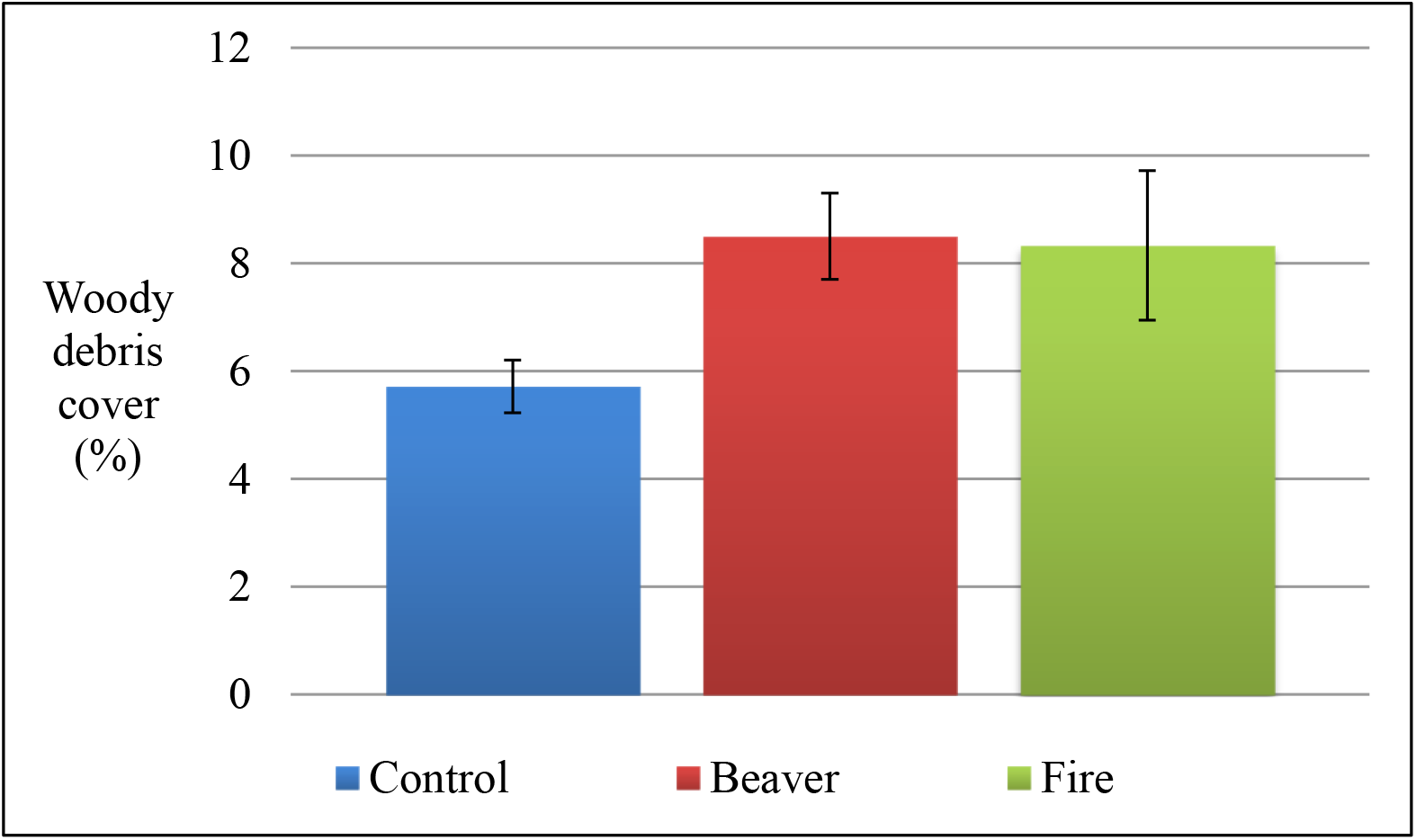
Mean (±standard error) density of woody debris at 0-3 meter height between control, beaver, and wildfire-influenced sites, San Pedro Riparian National Conservation Area, 2010.

## DISCUSSION

There was no significant difference in foliage density between control and beaver sites, but a significant difference existed between control and wildfire sites, and between beaver and wildfire sites. Lower foliage density was associated with wildfire sites rather than control or beaver sites. High-severity wildfire commonly kills cottonwood roots, as evidenced by the amount of large tree-fall consisting of branches and trunks of killed cottonwoods with no resprouts in wildfire areas. The higher mortality and lower resprouting rates of cottonwood after wildfire (Smith and Finch 2017), and the significantly higher Fremont cottonwood foliage density at control and beaver sites compared to wildfire sites, suggests the ecological adaptation of Fremont cottonwood to resprout subsequent to beaver herbivory. Cottonwood may resprout from the base (McGinley and Whitham 1985) and resprouting cottonwood occurs at beaver sites spanning a maximum of 10 years post-reintroduction. Conversely, wildfire sites spanned a range of 10 to 19 years since the wildfire occurred. Even with a longer time period since wildfire occurrence, past wildfire had more effect on cottonwood foliage density than beaver herbivory even given the longer time span since wildfire occurred.

Similarly, there was a significantly higher foliage density of Goodding’s willow at control and beaver sites compared to wildfire sites, but not a significant difference between control and beaver sites. Mean Goodding’s willow foliage density (approximately 50%) was higher than Fremont cottonwood (approximately 3%) at beaver influenced sites and higher (approximately 20%) than cottonwood (approximately 1%) at wildfire sites, indicating the superior ability of willow to resprout after beaver herbivory and wildfire. Old wildfire scars in the riparian area are apparent, demonstrating qualitatively that Goodding’s willow regrew after the wildfire and much of the Fremont cottonwood was removed.

Foliage density over all species between control, beaver, and wildfire sites was similar between 0 to 1 and 1 to 2-m (0 to 3.3 and 3.3 to 6.6-ft) in height, but density decreased significantly between 1 to 2 and 2 to 3-m (3.3 to 6.6 and 6.6 to 9.8-ft), and between 0 to 1 and 2 to 3-m (0 to 3.3 and 6.6 to 9.8-ft) in height. The interaction between density and height among control, beaver, and wildfire was near the significance level. Seep willow had significantly higher mean foliage density at control and beaver sites compared to wildfire sites, but not a significant difference between control and beaver sites. This may be attributed to thick stands of seep willow, reaching nearly 3-m (9.8-ft) in height over much of the San Pedro River’s edge. In contrast to Goodding’s willow and especially Fremont cottonwood, seep willow readily resprouts after removal by beaver or wildfire, quickly regaining its prior height and density in the matter of a few years. Thus, seep willow may account for the likeness in foliage density between control, wildfire, and beaver sites at 0 to 1-m (0 to 3.3-ft) height.

Similarly, significant differences in foliage density between control, beaver, and wildfire sites between the 1 to 2 and 2 to 3-m (3.3 to 6.6 and 6.6 to 9.8-ft) height may also be due to life history traits of vegetation species. Both Fremont cottonwood and Goodding’s willow grow to maximum heights taller than seep willow, with Goodding’s willow and Fremont cottonwood attaining approximately 20 and 30-m (66 and 98-ft) in height, respectively. As discussed previously, cottonwood did not resprout after high-severity wildfire, although Goodding’s willow readily resprouted, but takes longer to reach its maximum height than seep willow. These results indicate that the significant decrease in foliage density between 1 to 2 and 2 to 3-m (3.3 to 6.6 and 6.6 to 9.8-ft) may be from wildfire that removes Goodding’s willow and Fremont cottonwood, at least for longer time periods than beaver herbivory and longer times than seep willow from wildfire.

Large woody debris cover was not significantly different between control, beaver, and wildfire sites. The San Pedro River reaches flood events of several hundred cubic m/sec (thousands of cubic ft/sec) approximately every second or third year, with these flushing flows capable of moving woody debris throughout the main stem of the river. These flood events may redistribute woody debris so that any differences between control, beaver, and wildfire sites are negated.

Large woody debris is used by BLM as an indicator of riparian health during Proper Functioning Condition assessment (PFC; USDI Bureau of Land Management 1998). Indicator 13 of this assessment investigates whether floodplain and channel characteristics, such as rocks, overflow channels, coarse and/or large woody debris, are adequate to dissipate energy from flood flows. Wildfire may remove woody debris, while beaver may increase the amount of downed trees, at least locally and temporarily. Therefore, it is important that recruitment of younger tree stands from scouring natural flood regimes and redistribution of woody debris remain important ecohydrological processes of the San Pedro River.

Results indicate that Fremont cottonwood, Goodding’s willow, or seep willow do not recover similarly following beaver herbivory or wildfire events, even though each species is capable of resprouting under suitable environmental conditions. The 2012 National Riparian Service Team (NRST) PFC assessment for the SPRNCA notes that impacts of wildfire on cottonwood galleries was evident, but large-scale destabilization of banks or other adverse effects to the channel were not observed (USDI Bureau of Land Management 2012). Rather, wildfire promoted shrubs and herbaceous plants that also stabilize banks. Nevertheless, recommendations from the NRST included maintenance of the maximum number of cottonwood stands to achieve riparian function and associated resource values, because it is expected that cottonwood on the terraces will decrease in extent as trees become senescent. The risk of increasing wildfire size and severity is great given exotic species’ adaption to fire, fuel loading, drought, and climate warming (Smith and Finch 2017).

Perhaps these few years since reintroduction were still too early to ascertain quantifiable effects from beaver herbivory to the riparian vegetation parameter addressed during this study. However, beaver effects may become stronger with more time post-reintroduction. For example, cottonwood resprouts from beaver herbivory were relatively infrequent during 2008, with only a few observed, but appeared more frequent and with increased height during later years. Resprouts from beaver herbivory and/or seedling recruitment in beaver sites after flood events may eventually change foliage density between control and beaver sites in the future. Other future changes may include expanded wetlands created by beaver habitats, which may also reduce wildfire risk in these areas due to increased water, relative humidity, and fuel moisture.

## MANAGEMENT IMPLICATIONS FOR SPRNCA

Monitoring and maintenance of the beaver population should continue. Effects to ecohydrological components after beaver reintroduction are little-known, and assessments should continue.

Better wildfire planning and preparedness should be more fully integrated into land management planning to reduce the size and severity of future fire events, and their impacts to riparian and wetland habitats. Wildfire planning should include the following concepts.

Tamarisk did not occur within the sites used for this study, beaver herbivory to tamarisk was observed on SPRNCA on only one small plant, and tamarisk is not known to be a species favored by beaver. Because longer-term browsing by beaver may cause vegetation to be replaced by shrubs of non-preferred species (Donkor and Fryxell 2000), foliage density of tamarisk may change over time given beaver herbivory to preferred species. Tamarisk is highly adapted to wildfire, outcompeting native plants after wildfire. (Smith et al. 2009). With potential effects of beaver herbivory and wildfire to native species, and tamarisk’s superior adaptation to fire, tamarisk control should continue in order to manage for a possible community shift away from native species.

Prescribed fire on SPRNCA should be conducted in grassland and upland habitats in order to reduce fuel loading, create natural fuel breaks, and thereby protect the cottonwood/willow gallery forest from high-severity wildfire. Even so, prescribed fire would not likely prevent all wildfires from entering or starting in the riparian gallery, and a mosaic of habitat would be expected following wildfire. This mosaic may be important for future cottonwood recruitment.

---

**Table 1.** Mean Fremont cottonwood, Goodding’s willow, seep willow, and woody debris cover (in percent) at 0-1, 1-2, and 2-3 meters above ground between control, beaver, and wildfire-influenced sites, San Pedro Riparian National Conservation Area, 2010.

| Species/Height (m) | Control | Beaver | Wildfire |
| --- | --- | --- | --- |
| Fremont cottonwood |  |  |  |
| 0-1 | 55.8 | 42.3 | <1 |
| 1-2 | 51.7 | 50.0 | <1 |
| 2-3 | 54.6 | 52.3 | <1 |
| Goodding's willow |  |  |  |
| 0-1 | 46.8 | 39.1 | 8.1 |
| 1-2 | 50.2 | 50.0 | 19.4 |
| 2-3 | 42.5 | 51.4 | 33.1 |
| Seep willow |  |  |  |
| 0-1 | 63.3 | 58.9 | 49.1 |
| 1-2 | 57.3 | 54.3 | 49.2 |
| 2-3 | 45.0 | 46.9 | 30.6 |
| Woody debris |  |  |  |
| 0-1 | 8.6 | 11.5 | 16.7 |
| 1-2 | 3.6 | 10.5 | 3.9 |
| 2-3 | 5.0 | 3.5 | 4.5 |

## LITERATURE CITED

Bailey, Vernon. 1971. Mammals of the Southwestern United States. New York: Dover Publications, Inc. 412 p.

Bendix, Jacob Cowell, C. Mark. 2010. Impacts of wildfire on the composition and structure of riparian forests in Southern California. Ecosystems 13:99–107.

Busch, David E. 1995. Effects of fire on southwestern riparian plant community structure. The Southwestern Naturalist 40(3):259–267.

Donkor, Nobel T.; Fryxell, John M. 2000. Lowland boreal forests characterization in Algonquin Provincial Park relative to beaver (Castor canadensis) foraging and edaphic factors. Plant Ecology 148(1):1–12.

Johnson, Glenn E., and van Riper III, Charles. 2014. Effects of reintroduced beaver (Castor canadensis) on riparian bird community structure along the upper San Pedro River, southeastern Arizona and northern Sonora, Mexico. [Online]. U.S. Geological Survey Open-File Report 2014-1121, 98 p. Available: http://dx.doi.org/10.3133/ofr20141121.

Johnston, Carol A.; Naiman Robert J. 1990. Browse selection by beaver: effects on riparian forest composition. Canadian Journal of Forest Research 20(7):1036–1043.

McGinley, Mark A.; Whitham Thomas G. 1985. Central place foraging by beavers (Castor canadensis): a test of foraging predictions and the impact of selective feeding on the growth form of cottonwoods (Populus fremontii). Oecologia 66(4):558–562.

Sanders, S. D., and M. A. Flett. 1989. Ecology of a Sierra Nevada population of willow flycatchers (Empidonax traillii), 1986-1987. Administrative Report 88-3. Sacramento, CA: State of California, California Department of Fish and Wildlife. 27 p.

Smith, D. Max; Finch, Deborah, M. 2017. Climate change and wildfire effects in aridland riparian ecosystems: an examination of current and future conditions. General Technical Report RMRS-GTR-364. Fort Collins, CO: U.S. Department of Agriculture, Forest Service, Rocky Mountain Research Station. 65 p.

Smith, D. Max; Finch, Deborah M.; Gunning, Christian; [et al.]. 2009. Post-wildfire recovery of riparian vegetation during a period of water scarcity in the southwestern USA. Fire Ecology Special Issue 5(1):38–55.

Stromberg, Juliet; Tellman, Barbara, eds. 2009. Ecology and Conservation of the San Pedro River. Tucson: The University of Arizona Press. 529 p.

USDI Bureau of Land Management. 1998. Riparian area management: a user guide to assessing Proper Functioning Condition and the supporting science for lotic areas. Technical Reference 1737-15. Denver, CO: U.S. Department of the Interior, Bureau of Land Management. 127 p.

USDI Bureau of Land Management. 2012. Riparian conditions along the San Pedro River: potential natural communities and factors limiting their occurrence. Prineville, OR: U.S. Department of the Interior, Bureau of Land Management, National Riparian Service Team. 27 p.

Zar, Jerrold H. 1999. Biostatistical Analysis. Upper Saddle River: Prentice-Hall, Inc.663 p. + Appendices and Index.

